# Growth, yield, and quality of hot pepper (*Capsicum annuum* L.) as affected by the combined application of liquid bio-slurry and NPS fertilizer rates in Kucha, Gamo zone, south Ethiopia

**DOI:** 10.64898/2026.11.04.764160

**Authors:** Abrham Shumbulo Ganta, Yitbarek Abrham Biramo

## Abstract

Hot pepper (Capsicum annuum L.) is a key cash and nutritional crop in Ethiopia, yet its productivity remains constrained by inadequate and imbalanced soil nutrient management. Therefore, the field experiment was conducted in Kucha woreda, Gamo Zone, South Ethiopia, during the 2024/25 cropping season to investigate the effect of liquid bio-slurry and NPS fertilizer rates on growth, yield and quality of hot pepper. The experiment consisted of 16 treatments arranged in a factorial combination of four liquid bio-slurry (LBS) rates (0, 20, 40, and 60 m^3^ ha^−1^) and four NPS fertilizer rates (0, 75, 150, and 225 kg ha^−1^) using a randomized complete block design with three replications. All experimental data were collected following standard agronomic procedures and subjected to analysis of variance using GenStat statistical software (18^th^ edition). The results revealed that both the main and interaction effects of liquid bio-slurry and NPS significantly (p ≤ 0.01) influenced phenological, growth, yield, and quality parameters. The combined application of 150 kg ha^−1^ NPS and 60 m^3^ ha^−1^ LBS produced the highest marketable dry fruit yield (2.94 t ha^−1^) and total dry fruit yield (3.16 t ha^−1^), representing a 129.69% yield advantage over the control (1.28 t ha^−1^) in the case of marketable dry fruit yield. The findings demonstrate that integrated nutrient management using combined LBS and NPS fertilizers significantly enhanced the productivity and quality of hot pepper. Therefore, the application of 150 kg ha^−1^ NPS combined with 60 m^3^ ha^−1^ liquid bio-slurry is recommended for improved hot pepper production for the study area and similar agro-ecological conditions.

## 1. Introduction

Hot pepper (*Capsicum annuum* L.) is an important vegetable and spice crop widely cultivated in tropical and subtropical regions, where it plays a significant role in human nutrition, income generation, and food security. It belongs to the family Solanaceae, which includes tomato, potato, and eggplant, and ranks second only to tomato within the family Solanaceae [1]. Beyond its culinary value, hot pepper is rich in vitamins, antioxidants, and bioactive compounds, particularly capsaicinoids, which contribute to its nutritional and medicinal importance [2].

According to the [3] report, pepper production globally has expanded to 1,615,140 ha and 2,069,990 ha under dry and green pepper, with corresponding yields of 4,157,205 and 36,136,996 tons, respectively. In Ethiopia, however, productivity remains low, averaging 1.79 t ha^−1^ for dry pepper and 5.94 t ha^−1^ for green pepper [4], which is far below global potential. In the southern regions, including SNNPRS, yield performance is still constrained by multiple biophysical and management factors [5-7]. In the area, hot pepper is a key cash crop for smallholder farmers contributing significantly to household income and food security [8], and is widely grown in different forms such as sweet, chili, and pungent types [9].

Integrated nutrient management, which combines organic and inorganic nutrient sources, is widely recognized as a sustainable strategy for improving crop productivity, enhancing nutrient use efficiency, and maintaining soil fertility [10, 11]. Deficiencies of essential macronutrients such as nitrogen, phosphorus, and sulfur, as well as micronutrients including boron and zinc, are widely reported in Ethiopian soils [12]. Although mineral fertilizers improve yield due to their rapid nutrient availability, their continuous and sole application can lead to soil degradation, reduce nutrient use efficiency, and increase production costs [13]. In contrast, organic fertilizers such as compost, farmyard manure, and bio-slurry improve soil structure, microbial activity, and nutrient retention [14, 15]. Among these, bio-slurry, a byproduct of anaerobic digestion, is a valuable organic resource that supplies readily available nutrients, particularly ammoniacal nitrogen, and enhances soil fertility and biological activity [16, 17].

Previous studies have shown that combined applications of organic and mineral fertilizers improved crop performance more effectively [18, 19]. Similar findings have shown that organo-mineral fertilization systems significantly improve soil agrochemical properties and crop productivity compared to the application of only one type of nutrient, thus confirming the effectiveness of integrated nutrient management techniques [20]. However, adoption of integrated nutrient management practices in Ethiopia remains limited, as farmers predominantly rely on mineral fertilizers while crop residues are often removed for fuel and feed, accelerating soil fertility decline [21, 22].

In the study area, hot pepper production is constrained by poor soil fertility and inadequate nutrient management practices. Meanwhile, the newly introduced and increasing adoption of biogas technology in the area has improved the availability of bio-slurry, although its use in crop production remains very limited under farmers’ conditions. Farmers also rely heavily on blended fertilizers such as NPS without integrated nutrient strategies that could enhance efficiency and sustainability. Moreover, no research work has been done in the area for optimal utilization of integrated fertilizer combinations. Therefore, the combined use of liquid bio-slurry with NPS fertilizer is a promising approach for improving productivity while sustaining soil fertility. Despite these potential benefits, there is no empirical evidence on the combined effects of liquid bio-slurry and NPS fertilizer on the growth, yield, and quality of hot pepper under field conditions in Southern Ethiopia. Although some investigation was done elsewhere regarding integrated fertilizer use, as a matter of fact it does not work for the specific local environmental and edaphic conditions. Hence, the interaction effects of these nutrient sources on fruit yield, yield components, and quality traits have not been investigated for site-specific recommendations for sustainable hot pepper production.

Therefore, this study was conducted to evaluate the effects of the integrated application of liquid bio-slurry and NPS fertilizer on the growth, yield, and quality of hot pepper, and to determine the optimum combination of integrated fertilizer nutrient sources for sustainable production of hot pepper in the study area and similar agro-ecologies.

## 2. Materials and methods

### 2.1. Description of the study area

The field experiment was conducted at Kucha District, Gamo Zone, South Ethiopia. The geographical co-ordinates of the experimental site were 06°28′38.76″ N latitude, 37°28′3.37″ E longitude, and an altitude of 1356 m above sea level (Fig.1). The annual rainfall is 1350 mm, and the annual minimum and maximum temperatures were 21 °C and 25 °C, respectively (Fig.2). The soil is clay loam in texture and moderately acidic with a pH of 6.2.

**Fig. 1.**
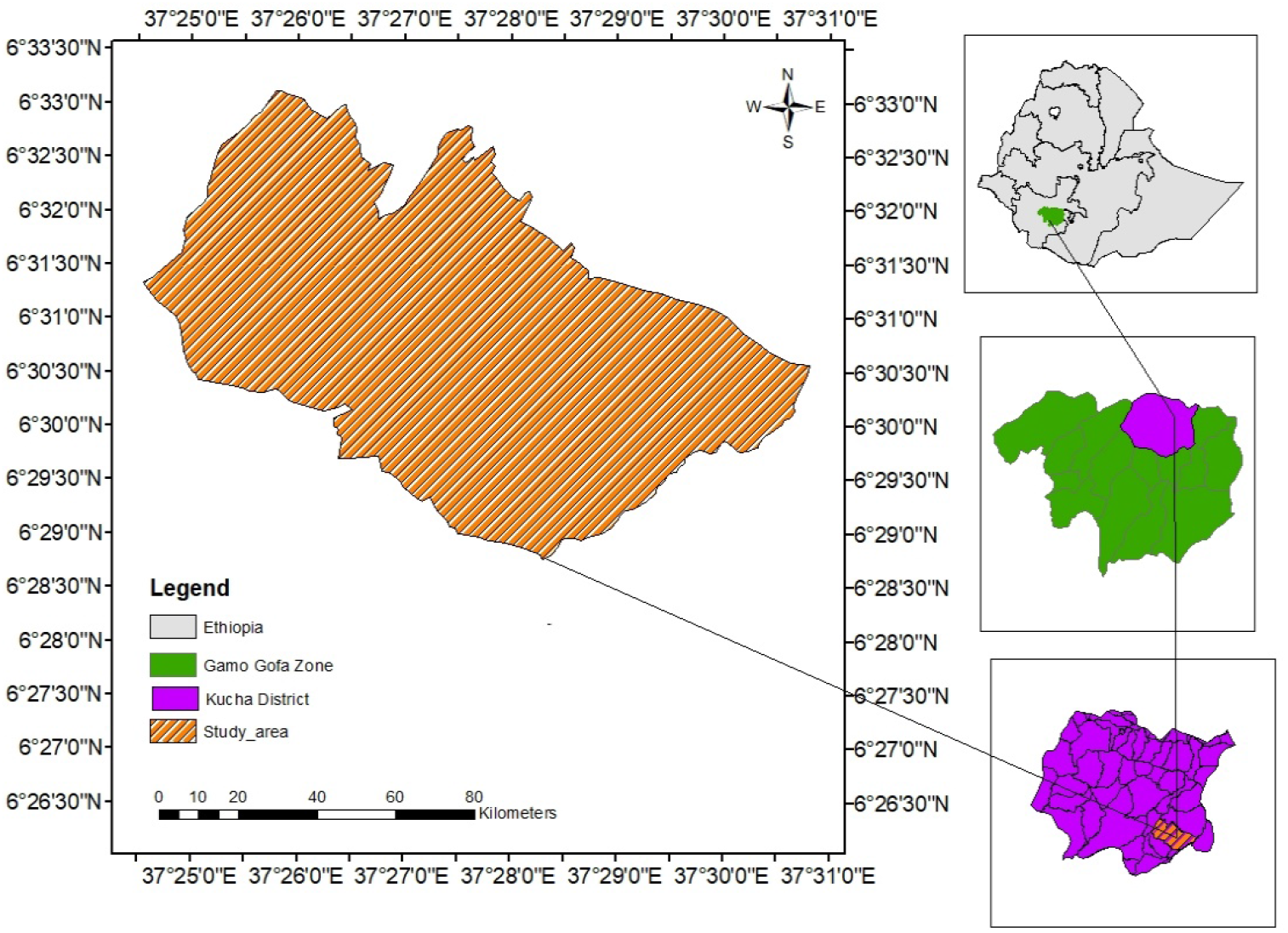
Map indicating the geographical location of the study area.

**Fig. 2.**
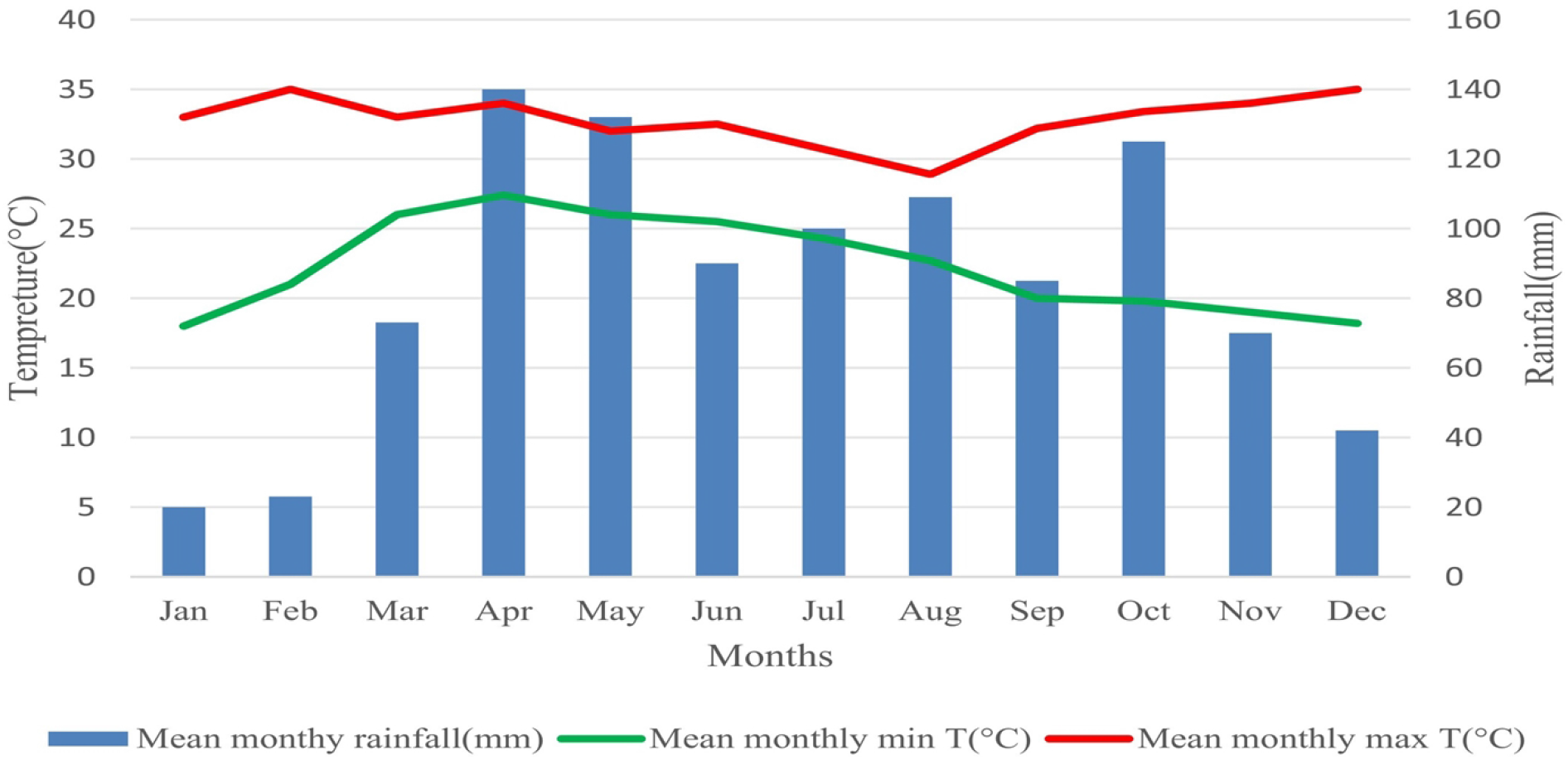
Average monthly annual rainfall along with minimum and maximum temperatures (1983– 2022) of the study area. Source: [23].

### 2.2. Description of experimental materials, treatments, design, and crop management

The experiment was conducted using well-fermented liquid bio-slurry, blended NPS and urea fertilizers, and the improved Mareko Fana hot pepper variety. The variety was recognized for its high yield, wide adaptability (1200–2000 m.a.s.l.), and desirable fruit quality traits, including large, pungent, dark-red pods with high powder yield [24, 25]. Treatments consisted of four rates of liquid bio-slurry (0, 20, 40, and 60 m^3^ ha^−1^) and four levels of NPS fertilizer (0, 75, 150, and 225 kg ha^−1^), arranged in a 4 × 4 factorial within a randomized complete block design with three replications. Each replication comprised 16 plots, 3.0 m x 3.2 m (9.6 m^2^), with 0.5 m and 1 m spacing between plots and blocks, respectively. Plants were spaced at 60 cm × 40 cm, giving 40 plants per plot, and data were collected from central rows to avoid border effects.

Seeds were obtained from Melkassa Agricultural Research Center, and seedlings were raised in a nursery under standard management practices, including mulching, irrigation, shading, and pest control. Seedlings were transplanted 45 days after sowing using healthy, uniform plants (20–25 cm height) following hardening. The measured NPS fertilizer and bio-slurry were applied at transplanting via side placement according to treatment levels, while urea (100 kg ha^−1^) was applied in split doses to enhance nitrogen use efficiency, with half applied at transplanting and the remaining half at the early vegetative stage with uniform agronomic activities. Supplementary irrigation was applied at 1–3 day intervals during establishment and 4–7 days thereafter, depending on environmental conditions [26]. Weed management and plant protection measures were carried out as required, including alternating applications of Megaban Plus (2 L ha^−1^) and Karate 5% EC (300 mL ha^−1^) for insect control and Mancozeb (3 kg ha^−1^) for fungal diseases. Fruits were harvested at physiological maturity and dried under partial shade to maintain quality. All practices followed recommended agronomic standards to ensure uniform crop performance and reliable treatment evaluation.

#### 2.2.1. Soil and liquid bio-slurry sampling and analysis

To assess the initial fertility status of the experimental site, soil samples were collected before transplanting from the 0–30 cm depth using a zigzag sampling method across the field. The samples were composited, and a representative subsample (1 kg) was air-dried, ground, and passed through a 2 mm sieve for analysis. Selected physicochemical properties were determined at the Sodo Soil Testing and Fertility Improvement Laboratory following standard procedures. Liquid bio-slurry samples were obtained from biogas digesters primarily fed with cow dung and locally available grasses. The slurry was thoroughly homogenized by gentle stirring to avoid sediment disturbance. Representative samples were collected from multiple digesters, composited, and a 1 L subsample was prepared for laboratory analysis. Key parameters analyzed included pH, organic carbon (OC), total nitrogen (TN), cation exchange capacity (CEC), available phosphorus (Av. P), and available sulfur (Av. S) using established analytical methods. Organic carbon was determined by the Walkley–Black wet oxidation method, total nitrogen by the micro-Kjeldahl method, pH in a 1:2.5 (sample: water) suspension, CEC by ammonium acetate extraction, and available phosphorus using the Olsen method.

The bio-slurry analysis revealed relatively high nutrient content, with 5.97% OC, 1.5% TN, 263 ppm available P, and 2.35 ppm available S, along with a slightly alkaline pH (7.5) and high CEC (60 cmol (+) kg^−1^). The moisture content was 94.1%, corresponding to 5.9% dry matter, indicating its suitability as a nutrient-rich organic amendment. The experimental soil was classified as clay loam with a slightly acidic pH (6.2). It contained 1.09% organic carbon and 0.09% total nitrogen, both within the medium range, but had relatively low organic matter content (1.9%). Available phosphorus (26 ppm) was rated as medium, whereas CEC (9.2 cmol (+) kg^−1^) was considered low (Table 1). Available sulfur was determined using monocalcium phosphate extraction. Overall, the soil exhibited moderate fertility constraints, particularly in organic matter and nutrient-holding capacity, highlighting the need for integrated application of organic and inorganic fertilizers to enhance soil fertility and optimize hot pepper productivity under the study conditions.

**Table 1.** Selected physical and chemical properties of the experimental site soil and liquid bio-slurry test before transplanting.

| Parameters | Soil Values | Liquid Bio-Slurry Value |
|---|---|---|
| pH (H<sub>2</sub>O) | 6.2 | 7.5 |
| Total nitrogen (%) | 0.09 | 1.5 |
| Organic carbon (%) | 1.09 | 5.97 |
| Organic matter (%) | 1.9 | 30.4 |
| Cation exchange capacity (meq/100 gm) | 9.2 | 60 |
| Ava. P (ppm) | 26 | 263 |
| Ava. S (ppm) | 8.25 | 2.35 |
| Moisture contents% | - | 94.1 |
| Dry matter% | - | 5.9 |
| Soil texture |  |  |
| Sand (%) | 32 |  |
| Silt (%) | 29 |  |
| Clay (%) | 39 |  |
| Textural class: clay loam |  |  |

**Table 1.** Mean square value for the main and interaction effects of LBS and NPS fertilizer on phenology, growth, yield, and quality parameters of hot pepper.

| Variables | Rep | LBS | NPS | LBS*NPS | Error | CV |
|---|---|---|---|---|---|---|
| Degree of freedom | 2 | 3 | 3 | 9 | 30 |  |
| Days to 50% flowering | 29.24 | 153.736 ** | 288.89 ** | 4.43 ** | 1.74 | 2.19 |
| Days to 50% fruit setting | 48.62 | 398.850 ** | 728.76 ** | 26.56 ** | 11.86 | 3.64 |
| Days to 90% maturity | 52.74 | 294.592 ** | 637.78 ** | 37.48 ** | 16.04 | 3.45 |
| Plant height | 69.49 | 613.598 ** | 449.49 ** | 9.26 ** | 3.64 | 3.15 |
| Number of branches/plant | 9.99 | 143.953 ** | 214.99 ** | 7.39 ** | 2.394 | 9.07 |
| Number of fruit/plant | 0.62 | 126.637 ** | 318.57 ** | 7.79 ** | 0.988 | 4.12 |
| Fruit length | 0.199 | 3.4178 ** | 10.21 ** | 0.39 ** | 0.0957 | 3.47 |
| Fruit width | 0.0125 | 0.55928 ** | 1.125 ** | 0.029** | 0.0092 | 4.14 |
| Marketable fruit yield | 0.035 | 1.24919 ** | 1.962 ** | 0.063 ** | 0.0128 | 5.03 |
| Unmarketable fruit yield | 0.00033 | 0.02140 ** | 0.016 ** | 0.0013** | 0.0002 | 7.29 |
| Total fruit yield | 0.00114 | 0.56477 ** | 2.003 ** | 0.0296** | 0.01 | 4.05 |
| Number of seeds/fruit | 69.686 | 288.877 ** | 828.68 ** | 128.65** | 20.56 | 4.76 |
| Total soluble solute | 0.00017 | 0.04549 ** | 0.116 *** | 0.97 *** | 0.00002 | 2.85 |

Where LBS = liquid bio-slurry, NPS = nitrogen, phosphorus, sulfur, CV = coefficient of variation, \*\*\* implies a significant level at (*p* ≤ 0.001), and \*\* implies a significant level at *p* ≤ 0.01), respectively.

#### 2.2.2. Data collection and measurements

Growth, phenological, and yield data of hot pepper were collected from the central three rows (net plot area: 3.78 m^2^) following standard procedures [27]. Measurements were taken as described below.

#### 2.2.3. Phenological and growth parameters

Days to 50% flowering (days): Number of days from transplanting to the stage when approximately 50% of plants in a plot initiated flowering. Days to 50% fruit set (days): Number of days from transplanting to the stage when 50% of plants in a plot developed at least one green fruit. Days to 90% fruit maturity (days): Number of days from transplanting to the stage when 90% of plants in a plot produced at least one physiologically mature fruit, characterized by firm texture, thick pericarp, and shiny, deep green appearance [24]. Plant height (cm): Measured from the soil surface to the apical growing point at 90% fruit maturity from ten randomly selected plants within the net plot area, and expressed as mean values [28]. Number of branches per plant: recorded from ten randomly selected plants in the net plot area at the third harvest, and averaged for analysis.

#### 2.2.4. Yield parameters of hot pepper

Number of fruits per plant (count): Determined by counting fruits from ten randomly selected and tagged plants within the net plot area at each harvest (first, second, and third), and expressed as mean values. Fruit width (cm): The average fruit width of ten randomly selected marketable fruits from the net plot area was measured at the middle of each fruit at each harvest, and the mean value was computed. Fruit length (cm): Measured from the tip to the basal end of ten randomly selected marketable fruits per plot at each harvest, and averaged. Marketable fruit yield (t ha^−1^): Fruits that were uniform in size, shape, and color, and free from mechanical damage, pests, diseases, and physiological disorders were classified as marketable. These fruits were harvested from the net plot area, dried, weighed at each harvest, and summed across harvests. The total was converted to a hectare basis and expressed in t ha^−1^ [29].

Unmarketable fruit yield (t ha^−1^): Fruits exhibiting defects such as damage, cracking, discoloration, blemishes, or physiological disorders due to biotic or abiotic stresses were considered unmarketable. These were collected, dried, weighed at each harvest, and expressed in t ha^−1^ which are unmarketable [30]. Total fruit yield (t ha^−1^): Calculated as the sum of marketable and unmarketable fruit yields from all harvests and expressed on a hectare basis. Number of seeds per fruit (count): Determined by counting seeds from ten randomly selected marketable fruits per plot at each harvest and expressed as mean values. Total soluble solids (TSS) were measured as °Brix using a portable refractometer. Ten fruits were randomly collected from five plants from each plot. Juice was extracted by macerating fresh pulp and filtering it through a clean muslin cloth. The refractometer was calibrated with distilled water before use. A few drops of juice were placed on the prism, and readings were taken in triplicate. The prism was rinsed with distilled water between samples. The mean value was recorded as the final TSS for statistical analysis [31].

#### 2.2.5. Data analysis

Collected data on growth and yield parameters were subjected to analysis of variance (ANOVA) using the General Linear Model procedure using GenStat statistical software (18th edition). Treatment means were separated using the least significant difference (LSD) test at the 5% probability level when significant differences were detected. Correlation analysis was performed to evaluate relationships among growth and yield parameters using Pearson’s correlation coefficients, following standard procedures [32].

## 3. Results and discussion

### 3.1. Effects of blended NPS fertilizer and liquid bio-slurry on phenology of hot pepper

#### 3.1.1. Days to 50% flowering

Days to 50% flowering of hot pepper were highly significantly (*p* < 0.01) affected by the main and interaction effects of NPS fertilizer and liquid bio-slurry, indicating a strong combined influence on crop phenology (Table 1). The earliest flowering (52 days) was recorded in the control treatment, whereas the longest duration (73 days) was observed under the combined application of 225 kg ha^−1^ NPS and 60 m^3^ ha^−1^ LBS, demonstrating that increasing nutrient supply prolonged the flowering period (Fig. 3).

**Fig. 3.**
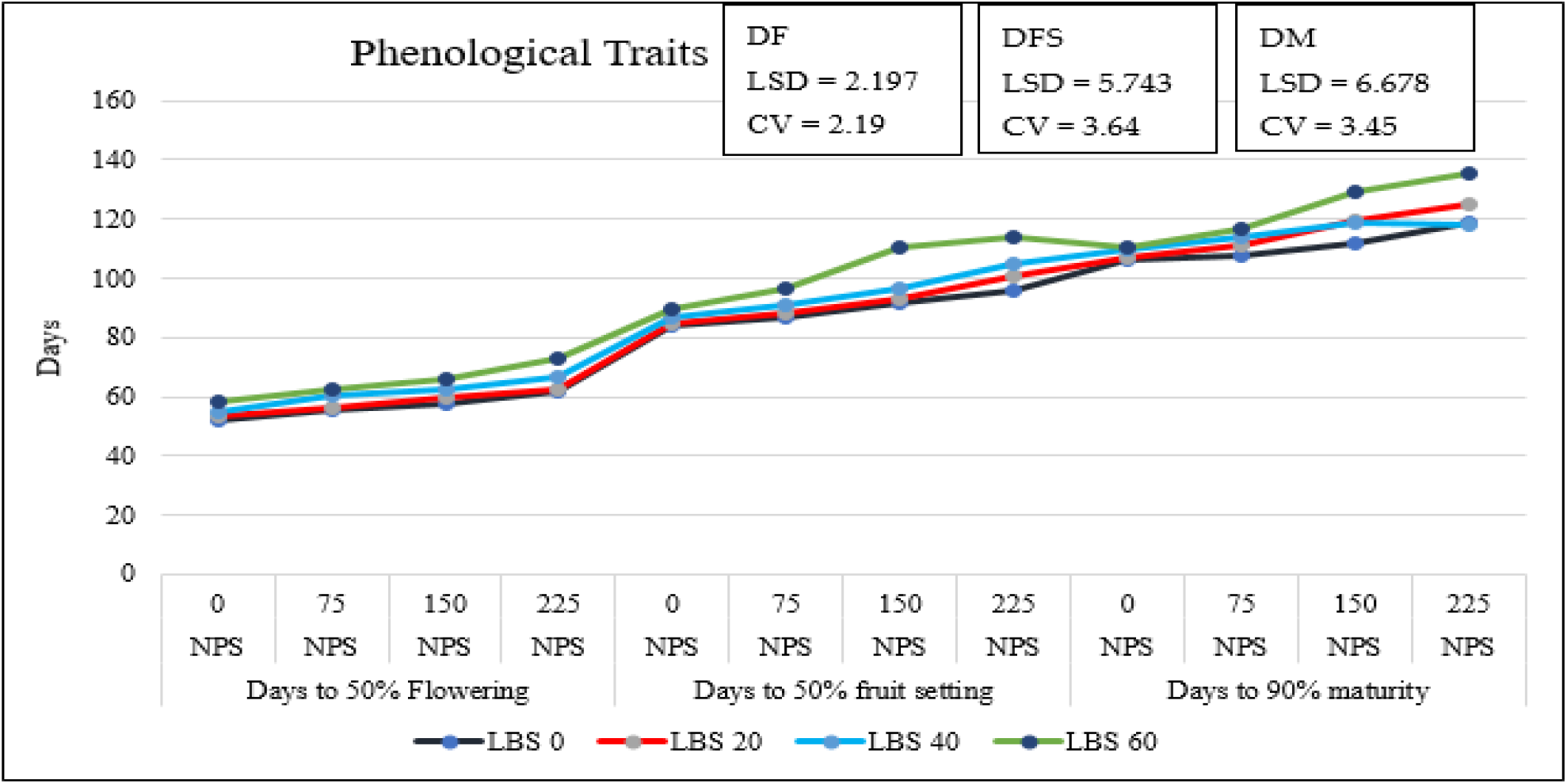
Interaction effects of liquid bio-slurry and NPS fertilizer on phenological parameters of hot pepper. Where LBS = liquid bio-slurry, NPS = nitrogen, phosphorus, Sulphur, CV = coefficient of variation, LSD = least significance difference, DF = days to 50% flowering, DFS = days to 50% fruit setting, DM = days to 90% maturity.

The delay in flowering under higher nutrient application can be attributed to enhanced vegetative growth driven by increased availability of nitrogen and phosphorus that extends the vegetative phase. The addition of liquid bio-slurry likely improved soil structure, microbial activity, and nutrient mineralization, further enhancing nutrient uptake and reinforcing vegetative growth. Consequently, the transition to the reproductive stage was delayed. In contrast, earlier flowering under low or no nutrient supply may be associated with nutrient stress, which accelerates reproductive development as a survival mechanism [33]. These findings were consistent with previous reports indicating that excessive nutrient availability, particularly nitrogen, promotes vegetative growth at the expense of reproductive development and delays flowering [34, 35].

#### 3.1.2. Days to 50% fruit setting

The current ANOVA result envisaged that the main effects of NPS fertilizer and liquid bio-slurry and their interaction effects significantly (*p* < 0.05) affected days to 50% fruit setting of hot pepper, indicating that integrated nutrient management has a significant influence on the phenology of the crop (Table 1). The earliest fruit setting (84.10 days) was recorded in the control treatment, whereas the longest duration (114.16 days) was observed under the combined application of 225 kg ha^−1^ NPS and 60 m^3^ ha^−1^ LBS, representing a delay of approximately 19 days compared to the control (Figure 3).

The delayed fruit setting under higher nutrient application can be attributed to higher rates of N obtained from bio-slurry, urea and NPS fertilizers that usually promote vigorous vegetative growth and prolong the transition from flowering to fruiting. Nitrogen-driven vegetative development, supported by phosphorus and sulfur-mediated metabolic processes, likely sustained continuous leaf and stem growth, thereby postponing reproductive development [36]. The contribution of liquid bio-slurry in improving soil conditions and nutrient mineralization further reinforces this effect [14]. In contrast, earlier fruit setting under low nutrient supply may result from nutrient stress, which accelerates reproductive processes. These findings are consistent with previous studies reporting that higher rates of organic and inorganic fertilizers delay fruit setting by extending vegetative growth, whereas lower rates promote earliness [37].

#### 3.1.3. Days to 90% fruit maturity

Days to 90% fruit maturity of hot pepper were significantly (*p* < 0.01) influenced by the main effects of NPS fertilizer and liquid bio-slurry, while their interaction showed a significant effect, indicating that integrated nutrient application markedly affected crop maturity (Table 1). The earliest maturity (106.10 days) was recorded in the control treatment, whereas the longest duration (135.43 days) was observed under the combined application of 225 kg ha^−1^ NPS and 60 m^3^ ha^−1^ LBS, followed by 150 kg ha^−1^ NPS with 60 m^3^ ha^−1^ LBS (129.50 days) (Fig. 3).

This is due to the fact that the extended maturity period is the consequence of delayed flowering in cases of higher nitrogen levels obtained from bio-slurry and NPS which promote vigorous vegetative growth, increased leaf area, and prolonged photosynthetic activity, thereby delaying the transition to physiological maturity. The contribution of liquid bio-slurry in improving soil properties and nutrient mineralization likely reinforced nutrient uptake and sustained assimilate production, further extending crop duration. In contrast, earlier maturity under nutrient-limited conditions may result from reduced vegetative growth, leading to a shortened life cycle and accelerated reproductive development. These findings were consistent with previous reports that indicated higher rates of nitrogen prolong vegetative growth and delay flowering, fruiting, and maturity in hot pepper [38 – 40].

### 3.2. Effect of blended NPS fertilizer and liquid bio-slurry on growth of hot pepper

#### 3.2.1. Plant height

Plant height of hot pepper was significantly (*p* < 0.05) influenced by the main effects of NPS fertilizer, liquid bio-slurry, and their interaction (Table 1). The tallest plant height (75 cm) was recorded by the combined application of 225 kg ha^−1^ NPS and 60 m^3^ ha^−1^ liquid bio-slurry, representing an increase of 30.8 cm compared to the control (44.2 cm), whereas the shortest plants were observed in unfertilized plots (Table 2).

**Table 2.**
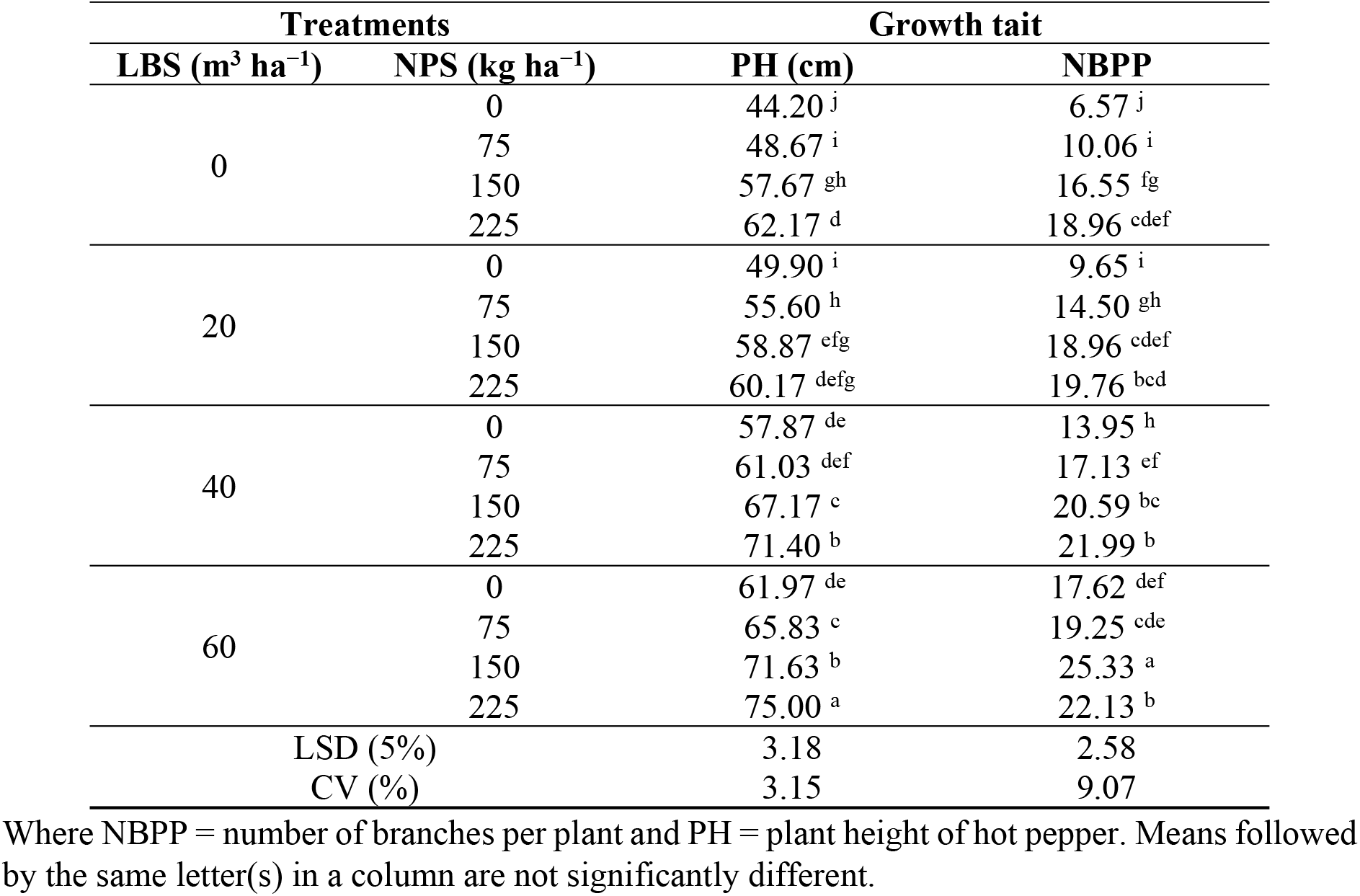
Interaction effects of liquid bio-slurry and NPS fertilizers on the number of branches and plant height of hot pepper.

The increase in plant height with higher nutrient rates can be attributed to enhanced photosynthesis and vigorous vegetative growth driven by nitrogen, phosphorus, and sulfur. Nitrogen promotes chlorophyll formation and shoot growth, phosphorus supports root and shoot development through high metabolic activity and cell division, and sulfur enhances nutrient availability and metabolic efficiency [12, 41]. Additionally, liquid bio-slurry improves soil structure, cation exchange capacity, water retention, and nutrient availability, further supporting stem elongation and biomass accumulation [42]. These results are consistent with previous studies showing that combined organic and inorganic fertilizer application enhances plant height through improved soil fertility and nutrient supply, which stimulates cell division and elongation [43, 44]. Similar trends were reported by [45 – 47], where the tallest pepper plants were obtained from integrated organic and mineral fertilizer treatments, while the shortest plants were observed under control conditions.

#### 3.2.2. Number of branches

The analysis of variance showed that the number of branches was significantly (*p* < 0.01) influenced by the main and interaction effects of NPS and liquid bio-slurry fertilizer rates (Table 1). The maximum number of branches (25.33) was recorded by application of both 150 kg ha^−1^ NPS and 60 m^3^ ha^−1^ liquid bio-slurry fertilizer, whereas plants without NPS and LBS recorded the lowest number of primary branches (3.65 and 3.85), respectively (Table 2).

The difference observed in the branching of the hot pepper plant at higher rates of NPS and LBS might be due to the availability of nutrients. The ample application of NPS leads to the production of more photosynthetic surface and subsequent accumulation of assimilates for more vegetative growth and ultimately gives a greater number of branches. Nitrogen is also pertinent in the metabolic activities of plants and photosynthetic activity, which affects the branching of pepper plants [48].

[49] reported that the maximum number of branches (10) was recorded at the highest level of NPK (90-60-30 kg ha^−1^), while the minimum number of branches (7.53) was recorded from the control treatment. Similarly, [48] reported that the maximum number of branches was recorded from the highest nitrogen level (150 kg ha^−1^), while the minimum number of branches was recorded by the control. According to [50], the maximum number of primary (3.33) and secondary (6.93) branches was recorded from the application of blended fertilizers at rates of 150 kg NPSB + 42 kg N ha^−1^ and 150 kg NPSBZn + 44 kg N ha^−1^, respectively.

In addition, liquid bio-slurry might be attributed to improved soil conditions (moisture retention, soil structure, aeration, and increased nitrogen availability), creating a suitable condition for plant growth and branching. In conformity with this result, [51] reported that application of poultry manure at 22 t ha^−1^ produced the highest number of branches, while the least number of branches per plant was recorded in the control treatment. Similarly, [51] reported that cow dung treatments at 7.5 t ha^−1^ gave the highest values of branch numbers (44), while the lowest number of branches (35.1) was recorded by the control. [52] Also reported that increasing the amount of poultry manure from 0 to 20 increased the number of branches of the pet plant from 2.5 to 7.

### 3.3. Effects of blended NPS fertilizers and liquid bio-slurry on the yield of hot pepper

#### 3.3.1. Number of fruits per plant

The analysis of variance showed that the main effects of NPS, liquid bio-slurry rates, and their interaction had a significant (*p* < 0.05) effect on the number of pods per plant of hot pepper (Table 1). Increasing the rate of NPS, as well as liquid bio-slurry, generally increased the number of pods per plant of pepper to a certain level. Regarding the interaction effect, the highest number of fruits per plant (33.76) was recorded by the combined application of 150 kg ha^−1^ NPS with 60 m^3^ ha^−1^ LBS, followed by the combined application of 225 kg ha^−1^ NPS with 40 m^3^ ha^−1,^ which resulted in 31.50 fruit numbers per plant. The lowest number of fruits per plant (12.92) was recorded by the control plot, which was statistically similar to treatment combinations of 75 kg ha^−1^ NPS with 0 m^3^ ha^−1^ LBS and 0 kg ha^−1^ NPS with 20 m^3^ ha^−1^ LBS with the number of fruits per plant being 17.27 and 18.73 respectively (Table 3).

**Table 3.** Interaction effects of liquid bio-slurry and blended NPS fertilizers on the number of fruits per plant, fruit length, and fruit width of hot pepper.

| **Treatments** |  | **Yield parameters** |  |  |  |
|---|---|---|---|---|---|
| **LBS (m<sup>3</sup> ha<sup>−1</sup>)** | **NPS (kg ha<sup>−1</sup>)** | **NFPP** | **FL (cm)** | **FW (cm)** | **NSPF (Count)** |
| 0 | 0 | 12.92<sup>i</sup> | 6.79<sup>g</sup> | 1.66<sup>j</sup> | 83.43<sup>g</sup> |
|  | 75 | 17.26<sup>h</sup> | 7.77<sup>ef</sup> | 1.75<sup>ij</sup> | 88.55<sup>efg</sup> |
|  | 150 | 23.34<sup>ef</sup> | 9.32<sup>c</sup> | 2.28<sup>fg</sup> | 92.83<sup>def</sup> |
|  | 225 | 26.22<sup>d</sup> | 9.54<sup>bc</sup> | 2.35<sup>efg</sup> | 96.45<sup>bcd</sup> |
| 20 | 0 | 18.73<sup>gh</sup> | 7.53<sup>f</sup> | 1.85<sup>hi</sup> | 85.74<sup>fg</sup> |
|  | 75 | 20.00<sup>g</sup> | 8.19<sup>e</sup> | 2.23<sup>g</sup> | 88.67<sup>efg</sup> |
|  | 150 | 26.30<sup>d</sup> | 9.18<sup>cd</sup> | 2.51<sup>bcd</sup> | 93.15<sup>def</sup> |
|  | 225 | 28.64<sup>c</sup> | 9.57<sup>bc</sup> | 2.59<sup>bc</sup> | 101.01<sup>bc</sup> |
| 40 | 0 | 19.86<sup>g</sup> | 8.03<sup>ef</sup> | 1.96<sup>h</sup> | 87.36<sup>efg</sup> |
|  | 75 | 21.99<sup>f</sup> | 8.76<sup>d</sup> | 2.46<sup>cde</sup> | 90.37<sup>defg</sup> |
|  | 150 | 28.51<sup>c</sup> | 9.62<sup>bc</sup> | 2.65<sup>ab</sup> | 96.29<sup>bcd</sup> |
|  | 225 | 31.50<sup>b</sup> | 9.87<sup>b</sup> | 2.64<sup>ab</sup> | 116.25<sup>a</sup> |
| 60 | 0 | 22.40<sup>f</sup> | 8.74<sup>d</sup> | 2.19<sup>g</sup> | 88.27<sup>efg</sup> |
|  | 75 | 24.71<sup>de</sup> | 9.39<sup>bc</sup> | 2.41<sup>def</sup> | 94.73<sup>cde</sup> |
|  | 150 | 33.74<sup>a</sup> | 10.48<sup>a</sup> | 2.76<sup>a</sup> | 117.55<sup>a</sup> |
|  | 225 | 29.40<sup>c</sup> | 9.68<sup>bc</sup> | 2.63<sup>ab</sup> | 103.37<sup>b</sup> |
| LSD (5%) |  | 1.66 | 0.52 | 0.16 | 7.5680 |
| CV (%) |  | 4.12 | 3.47 | 4.14 | 4.76 |

Where NFPP = number of fruit per plant, FL = fruit length, FW = fruit width, and NSPF = number of seeds per fruit. Means followed by the same letter(s) in a column are not significantly different.

The lowest number of fruits per plant recorded in the unfertilized treatment could be attributed to inadequate nutrient availability. In contrast, the number of fruits per plant increased significantly with the balanced and combined application of LBS and NPS blended fertilizers. This increment might be associated with higher levels of essential growth nutrients and soil moisture, which promote flower initiation and development, thereby increasing fruit set per plant. Furthermore, the application of LBS likely enhanced nutrient solubilization and availability, leading to improved uptake of NPS fertilizers and, consequently, higher fruit production.

The present findings were consistent with those of [53], who reported that the combined application of 75 kg ha^−1^ N, 25 kg ha^−1^ P_2_O_5_, and 7.5 t ha^−1^ farmyard manure resulted in a significant increase in pod number per plant. Similarly, [54] observed that the maximum fruit set (4.77) was achieved by the combined application of the recommended nitrogen rate and 50% vermicompost ha^−1^, whereas the minimum fruit set (2.40) was recorded in the control treatment. In agreement with the current study, [55] reported that the highest fruit set percentage (45.66%) was obtained with the application of 50% FYM combined with 50% urea, while the lowest fruit set (29.78%) was observed with 100% vermicompost alone. Moreover, [56] found that plants treated with 75% of the recommended NP fertilizer combined with 11.5 t ha^−1^ vermicompost produced 89.94% more fruits per plant compared to the unfertilized control. Likewise, [57] reported that the integrated application of 150 kg ha^−1^ NPS and 2.5 t ha^−1^ organic fertilizer significantly increased pod number per plant by 227% over the control.

#### 3.3.2. Fruit length

The analysis of variance indicated that fruit length of hot pepper was highly significantly (*p* ≤ 0.01) influenced by NPS and liquid bio-slurry application rates, as well as their interaction (Table A1). Fruit length generally increased with increasing rates of NPS and LBS up to an optimum level, beyond which no significant improvement was observed. The maximum fruit length (10.47 cm) was obtained from the combined application of 150 kg ha^−1^ NPS and 60 m^3^ ha^−1^ LBS, followed by 225 kg ha^−1^ NPS with 40 m^3^ ha^−1^ LBS. In contrast, the minimum fruit length (6.79 cm) was recorded in the control treatment, which was statistically similar to 0 kg ha^−1^ NPS + 20 m^3^ ha^−1^ LBS and 75 kg ha^−1^ NPS + 0 m^3^ ha^−1^ LBS (Table 3).

The observed increase in fruit length under integrated nutrient applications may be attributed to enhanced photosynthetic activity and assimilate production due to higher-level application of nutrients through NPS and LBS. These assimilates are subsequently translocated from the source (leaves) to the sink (developing fruits), promoting fruit growth. Optimal fertilization enhances biomass production and assimilate partitioning, resulting in increased fruit size, including length [58]. These findings were consistent with previous reports. [59] Noted that adequate nutrient supply improves vegetative growth, leaf area, and photosynthetic efficiency, leading to better assimilate partitioning and fruit development. Similarly, [50] highlighted the importance of balanced nutrient application from both organic and inorganic sources in improving fruit size. [55] Also reported increased fruit length under integrated fertilization, particularly with combined applications of organic and inorganic nutrient sources.

#### 3.3.3. Fruit width

The analysis of variance result showed that the fruit width of hot pepper was highly significantly (*p* ≤ 0.01) influenced by NPS and liquid bio-slurry application rates and their interaction (Table 1). Fruit width increased with increasing rates of NPS and LBS up to an optimum level. The maximum fruit width (2.76 cm) was recorded from the combined application of 150 kg ha^−1^ NPS and 60 m^3^ ha^−1^ LBS, followed by 225 kg ha^−1^ NPS with 40 m^3^ ha^−1^ LBS (2.64 cm), with both treatments being statistically similar. The minimum fruit width (1.66 cm) was observed in the control treatment (Table 3).

The increase in fruit diameter under integrated nutrient applications may be attributed to better nutrient availability and uptake, which enhanced vegetative growth and assimilate production. This promotes efficient carbohydrate accumulation and translocation to developing fruits, resulting in larger fruit size. In addition, the combined use of organic and inorganic fertilizers likely improves soil organic matter and microbial activity, further enhancing nutrient use efficiency and crop performance.

These results were consistent with earlier findings. [60] Indicated that adequate nutrient availability contributes to increased fruit size due to enhanced assimilate accumulation. [48] Also reported that improved nutrient uptake is directly associated with better fruit development and size. Similarly, [59, 30], and [28] demonstrated that increasing nitrogen and phosphorus application significantly enhanced pod diameter. [55] Further confirmed that the integrated application of organic and inorganic fertilizers improves fruit diameter and overall yield components.

#### 3.3.4. Seed number per fruit

The analysis of variance showed that the main effects of liquid bio-slurry, NPS fertilizers, and their interaction were significant (*p* ≤ 0.01) for the number of seeds per fruit (Table 1). The highest number of seeds per fruit (117.55) was recorded by the combined application of 150 kg ha^−1^ NPS and 60 m^3^ ha^−1^ LBS, followed by 225 kg ha^−1^ NPS with 40 m^3^ ha^−1^ LBS (116.25), with no significant difference between these treatments. These combinations increased seed number per fruit by 29.03% and 28.24%, respectively, as compared to the control (83.43), which recorded the lowest value (Table 3).

The increase in seed number per fruit under integrated nutrient application may be attributed to enhanced photosynthetic capacity and assimilate partitioning, leading to improved fruit size (length and width) and, consequently, greater seed development. Seed number is closely associated with fruit size, as larger fruits tend to accommodate more seeds. These findings agree with previous reports. [61] Reported a positive correlation between fruit weight and seed number, while [50] highlighted that adequate nutrient supply enhances photosynthetic activity and assimilate distribution, thereby increasing both fruit number and seed set.

#### 3.3.5. Marketable fruit yield

The ANOVA result indicated that the main and the interaction effects of NPS and liquid bio-slurry significantly (*p* ≤ 0.01) affected marketable dry fruit yield of hot pepper (Table 1). The highest marketable dry fruit yield (2.94 t ha^−1^) was obtained from the combined application of 150 kg ha^−1^ NPS and 60 m^3^ ha^−1^ LBS, followed by 150 kg ha^−1^ NPS with 40 m^3^ ha^−1^ LBS (2.85 t ha^−1^), with both treatments being statistically similar. The maximum yield advantage of 129.69% was observed as compared to the control (1.28 t ha^−1^) (Figure 4).

**Figure 4.**
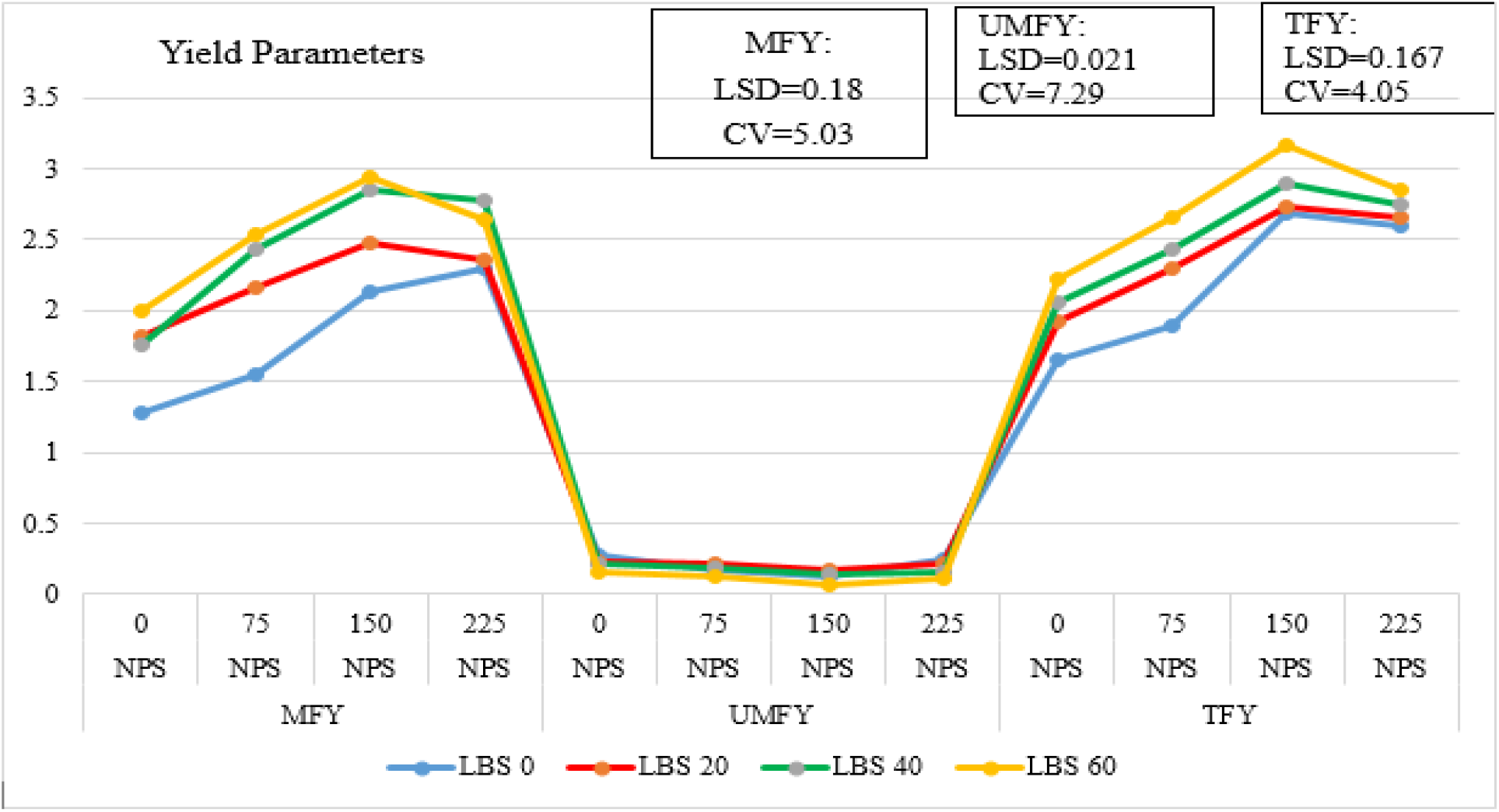
Interaction effects of liquid bio-slurry and blended NPS fertilizers on yield and yield-related parameters of hot pepper. Where MFY = marketable fruit yield, UMFY = unmarketable fruit yield, and TFY = total fruit yield.

The superior yield under integrated nutrient application may be attributed to balanced and sustained nutrient supply, synergetic effects of N, P, and S nutrients which enhanced vegetative growth, fruit development, and overall productivity. Improvements in yield components such as fruit length, fruit width, seed number, and fruit number per plant likely contributed to the observed increases in marketable yield which is a polygenic trait.

This confirmed the previous reports on the increased efficiency of nutrient acquisition and reduced stress effects on plants due to the application of organo-mineral fertilization, which contributes to the improvement of crop productivity and quality [62]. Consistent with the present study, [63] reported that the combined application of organic and inorganic fertilizers improves nutrient availability and crop performance, particularly in nutrient-deficient soils. Similarly, [55] demonstrated that integrated nutrient management results in higher marketable yields compared to sole applications of either organic or inorganic fertilizers.

#### 3.3.6. Unmarketable fruit yield

The analysis of variance revealed that the main effects of NPS and liquid bio-slurry were highly significant (*p* ≤ 0.01), while their interaction was also significant for the unmarketable dry fruit yield of hot pepper (Table 1). The highest unmarketable yield (0.27%) was recorded in the control treatment, whereas the lowest value (0.07%) was obtained from the combined application of 150 kg ha^−1^ NPS and 60 m^3^ ha^−1^ LBS (Figure 4).

The reduced unmarketable yield under integrated nutrient application may be attributed to balanced nutrient supply, which enhances fruit development and increases the proportion of marketable fruits. In contrast, higher unmarketable yield in the control and low-input treatments is likely due to nutrient deficiencies that impair physiological processes and fruit development.

These findings were consistent with previous reports of [50], who observed that low phosphorus levels result in smaller, less firm fruits with higher defect incidence. Nitrogen, for instance, is a key component of chlorophyll, enzymes, and other biomolecules essential for plant growth [64], and its deficiency can limit productivity and fruit quality. Similarly, an inadequate phosphorus supply may restrict root development and assimilate translocation, leading to malformed, undersized, or defective fruits. Likewise, [50] reported the highest unmarketable yield in unfertilized plots and the lowest under optimal integrated fertilization. Overall, insufficient or imbalanced nutrient supply contributes to increased unmarketable yield through the production of underdeveloped and poor-quality fruits.

#### 3.3.7. Total fruit yield

The analysis of variance revealed that the total dry fruit yield of hot pepper was significantly (*p* ≤ 0.01) affected by NPS and liquid bio-slurry application rates, while their interaction was also significant (Table 1). Total yield increased with increasing rates of both NPS and LBS up to an optimum level. The highest total dry fruit yield (3.16 t ha^−1^) was obtained from the combined application of 150 kg ha^−1^ NPS and 60 m^3^ ha^−1^ LBS, followed by 150 kg ha^−1^ NPS with 40 m^3^ ha^−1^ LBS (2.90 t ha^−1^). In contrast, the lowest yield (1.66 t ha^−1^) was recorded in the control. It was observed that a 90.36% total dry fruit yield increment was recorded as compared to the control (Figure 4). However, a decline in yield was observed at the highest fertilizer combination (225 kg ha^−1^ NPS with 60 m^3^ ha^−1^ LBS), indicating that yield response peaks at an optimal nutrient level.

The enhanced yield under integrated nutrient application can be attributed to improved soil fertility and nutrient supply, which promote plant growth, photosynthetic efficiency, and assimilate production. Nitrogen contributes to chlorophyll, protein, and enzyme synthesis, while phosphorus supports root development and fruit formation. In addition, organic inputs improve soil structure, microbial activity, and nutrient availability, thereby enhancing nutrient use efficiency and crop productivity.

These results were consistent with previous studies. [48] Reported significant improvements in pepper yield with nitrogen application due to its role in essential plant metabolic processes. [65] Highlighted the role of sulfur-containing fertilizers in improving nutrient availability through soil pH modification. Similarly, [57, 66], and [67] demonstrated that integrated use of organic and inorganic fertilizers enhances nutrient release, uptake, and yield sustainability. Other studies [68] also confirmed that combined nutrient management improves yield and yield components. High nitrogen levels, for instance, can promote excessive vegetative growth at the expense of reproductive development, thereby reducing yield. This trend is supported by [28, 69], and [70], who reported yield declines beyond optimal fertilizer rates.

#### 3.3.8. Total soluble solids of hot pepper

Total soluble solids (TSS) of hot pepper were significantly (*p* < 0.001) influenced by the main and interaction effects of NPS fertilizer and liquid bio-slurry, indicating a strong synergistic effect of integrated nutrient management (Table 1). The highest TSS (0.28 °Brix) was recorded from the combined application of 225 kg NPS ha^−1^ and 60 m^3^ liquid bio-slurry ha^−1^, whereas the lowest TSS (0.06 °Brix) was obtained from 75 kg NPS ha^−1^ combined with 20 m^3^ liquid bio-slurry ha^−1^ (Fig. 5). The increased TSS under higher combined nutrient application could be attributed to improved nutrient availability and uptake, which enhanced photosynthetic activity and carbohydrate accumulation in fruits. Nitrogen, phosphorus, and sulfur supplied from NPS are important for chlorophyll formation, energy transfer, and assimilate production, while liquid bio-slurry improves soil microbial activity and nutrient use efficiency. In contrast, lower nutrient applications reduced assimilation and sugar accumulation in fruits. This finding agrees with [71], who reported that improved nutrient availability and rhizosphere microbial activity enhance plant metabolic processes and fruit quality attributes, including soluble solids.

**Fig. 5.**
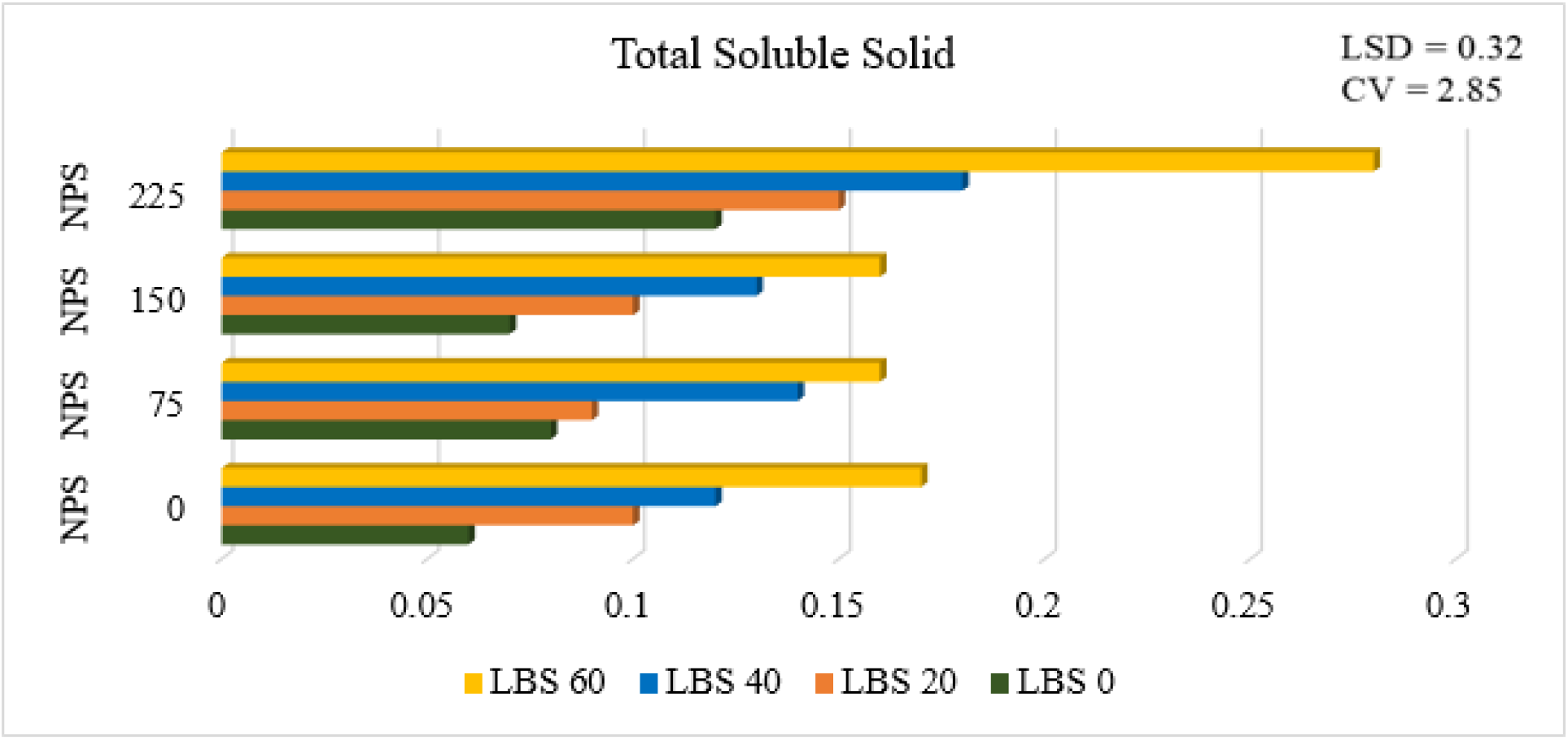
Interaction effects of NPS fertilizer and liquid bio-slurry on total soluble solids of hot pepper fruits

### 3.4. Association of phenology, growth, and yield parameters as influenced by NPS and LBS fertilizer rates

Correlation coefficient (r) values computed to determine the association between phenology, growth, and yield-related parameters are depicted in Table 4. The correlation values explained the apparent association of the phenology, growth, and yield parameters, indicating the magnitude and directions of the association and relationships.

**Table 4.** Correlation coefficient among different growth and yield parameters of hot pepper.

| **PAR** | **DF** | **DFS** | **DM** | **PH** | **NBPP** | **NFPP** | **NSPF** | **FL** | **FW** | **MFY** | **UMFY** | **TFY** |
|---|---|---|---|---|---|---|---|---|---|---|---|---|
| DF | 1 |  |  |  |  |  |  |  |  |  |  |  |
| DFS | 0.84 ** | 1 |  |  |  |  |  |  |  |  |  |  |
| DM | 0.80 ** | 0.91 ** | 1 |  |  |  |  |  |  |  |  |  |
| PH | 0.88 ** | 0.81 ** | 0.77 ** | 1 |  |  |  |  |  |  |  |  |
| NBPP | 0.82 ** | 0.80 ** | 0.75 ** | 0.90 ** | 1 |  |  |  |  |  |  |  |
| NFPP | 0.84 ** | 0.83 ** | 0.77 ** | 0.86 ** | 0.92 ** | 1 |  |  |  |  |  |  |
| NSPF | 0.67 ** | 0.82 ** | 0.69 ** | 0.70 ** | 0.73 ** | 0.82 ** | 1 |  |  |  |  |  |
| FL | 0.81 ** | 0.76 ** | 0.74 ** | 0.86 ** | 0.92 ** | 0.92 ** | 0.74 ** | 1 |  |  |  |  |
| FW | 0.79 ** | 0.76 ** | 0.73 ** | 0.83 ** | 0.92 ** | 0.89 ** | 0.72 ** | 0.87 ** | 1 |  |  |  |
| MFY | 0.80 ** | 0.75 ** | 0.72 ** | 0.86 ** | 0.89 ** | 0.90 ** | 0.73 ** | 0.86 ** | 0.93 ** | 1 |  |  |
| UMFY | −0.62 ** | −0.61 ** | −0.52 ** | −0.69 ** | −0.69 ** | −0.65 ** | −0.55 ** | −0.68 ** | −0.61 ** | −0.66 ** | 1 |  |
| TFY | 0.78 ** | 0.78 ** | 0.77 ** | 0.82 ** | 0.89 ** | 0.92 ** | 0.75 ** | 0.91 ** | 0.90 ** | 0.93 ** | −0.7 ** | 1 |

Where DF = days to 50% flowering, DFS = days to 50% fruit setting, DM = days to 90% fruit maturity, PH = plant height, NBPP = number of branches per plant, NFPP = number of fruit per plant, NSPF = number of seeds per fruit, FL = fruit length, FW = fruit width, MFY = marketable fruit yield, UMFY = unmarketable fruit yield, TFY = total fruit yield \*\*highly significant.

The correlation analysis showed that marketable fruit yield was highly significantly and positively correlated with days to 50% flowering (r = 0.80 **), days to 50% fruit setting (r = 0.75 **), days to maturity (0.72 **), plant height (r = 0.86 **), number of branches per plant (r = 0.89**), number of fruits per plant (r = 0.90 **), fruit length (r = 0.86 **), fruit width (r = 0.93 **), total fruit yield (r = 0.93 **) and number of seeds per fruit (r = .0.73**). Moreover, marketable fruit yield was significantly and negatively correlated with unmarketable fruit yield (r = −0.66 **) of pepper. Similarly, unmarketable fruit yield of hot pepper was highly significantly (*p* ≤ 0.01) and negatively correlated with fruit length (r =−0.68 **), fruit width (r = −0.61 **), number of seeds per fruit (r = −0.55 **), and total fruit yield (r =−0.70 **) (Table 4).

Positively correlated parameters showed that the yield was increased as any improvement in positively correlated parameters of pepper, such as number of fruits per plant, fruit length, fruit width, and seed number per fruit, while negatively correlated parameters showed that yield increased as parameters decreased.

## 4. Conclusions

The results of this study confirm that the integrated application of liquid bio-slurry and NPS fertilizer significantly enhances the growth, yield, and quality of hot pepper. Both main and interaction effects were highly significant across all parameters, demonstrating the effectiveness of integrated nutrient management. Although higher nutrient rates promoted vegetative growth and delayed phenological development, optimum yield was achieved at moderate application levels. The combined application of 150 kg ha^−1^ NPS with 60 m^3^ ha^−1^ LBS produced the highest marketable dry fruit yield (2.94 t ha^−1^) and total dry fruit yield (3.16 t ha^−1^), along with improved yield components and reduced unmarketable yield. In contrast, excessive nutrient application led to diminished yield performance, indicating the importance of balanced fertilization. Overall, integrating organic and inorganic nutrient sources proved superior to sole applications, with 150 kg ha^−1^ NPS with 60 m^3^ ha^−1^ LBS identified as the optimal combination for maximizing productivity and fruit quality while supporting sustainable soil fertility management.

## 5. Recommendation

Based on the findings of this study, the combined application of 150 kg ha^−1^ NPS and 60 m^3^ ha^−1^ liquid bio-slurry is recommended for hot pepper production in Kucha District and areas with similar agro-ecological conditions, as it maximizes yield, improves fruit quality, and reduces unmarketable yield. This integrated nutrient management approach offers a practical strategy to enhance productivity while improving soil fertility and nutrient use efficiency. Farmers are encouraged to utilize locally available bio-slurry in combination with mineral fertilizers to reduce reliance on costly inorganic fertilizers and promote sustainable crop production. However, excessive fertilizer application should be avoided, as higher rates did not result in additional yield benefits and may lead to nutrient imbalance. Future research should also consider economic analysis under multiple farming conditions and assess long-term soil health impacts of integrated nutrient management practices.

## Author Contributions

Abrham Shumbulo, the author, conceptualized the study, developed the methodology, conducted the field experiment, contributed to data collection & interpretation, and critically revised the manuscript for important intellectual content and journal publication.

Yitbarek Abrham, Co-author, contributed for project proposal development, field works, collected and organized the data, performed statistical analysis, and prepared the first draft of the manuscript.

## Data Availability Statement

All data generated or analyzed during this study are included in this manuscript.

